# Inter-individual variability in motor learning due to differences in effective learning rates between generalist and specialist memory stores

**DOI:** 10.1101/2024.07.15.603502

**Authors:** Rieko Osu, Norikazu Sugimoto, Toshinori Yoshioka, Nicolas Schweighofer, Mitsuo Kawato

## Abstract

Humans exhibit large interindividual differences in motor learning ability. However, most previous studies have examined properties common across populations, with less emphasis on interindividual differences. We hypothesized here, based on our previous experimental and computational motor adaptation studies, that individual differences in effective learning rates between a generalist memory module that assumes environmental continuity and specialist modules that are responsive to trial-by-trial environmental changes could explain both large population-wise and individual-wise differences in dual tasks adaptation under block and random schedules. Participants adapted to two opposing force fields, either sequentially with alternating training blocks or simultaneously with random sequences. As previously reported, in the block training schedule, all participants adapted to the force field presented in a block but showed large interference in the subsequent opposing force field blocks, such that adapting to the two force fields was impossible. In contrast, in the random training schedule, participants could adapt to the two conflicting tasks simultaneously as a group; however, large interindividual variability was observed. A modified MOSAIC computational model of motor learning equipped with one generalist module and two specialist modules explained the observed behavior and variability for wide parameter ranges: when the predictions errors were large and consistent as in block schedules, the generalist module was selected to adapt quickly. In contrast, the specialist modules were selected when they more accurately predicted the changing environment than the generalist, as during random schedules; this resulted in consolidated memory specialized to each environment, but only when the ratio of learning rates of the generalist to specialists was relatively small. This dynamic selection process plays a crucial role in explaining the individual differences observed in motor learning abilities.

## Introduction

Acquiring dexterous motor skills such as playing sports or musical instruments can take years of practice, although considerable individual variability exists (Ackerman & Cianciolo, 2000; Golenia et al., 2014; Stark-Inbar et al., 2017). This variability has largely been overlooked in quantitative and computational motor control studies, which typically only investigate population averages. Studies of motor adaptation with a single environment, i.e., repeated exposure to the same task, show consistent results with small individual variability, e.g., (Krakauer et al., 2005). In contrast, studies of adaptation to multiple tasks show inconsistent results; some studies have reported that humans can learn conflicting multiple tasks simultaneously (Forano & Franklin, 2020; Lee & Schweighofer, 2009; Osu et al., 2004; Shelhamer et al., 2005; Wada et al., 2003), while others indicate that such learning is difficult or even impossible (Gandolfo et al., 1996; Gupta & Ashe, 2007; Hinder et al., 2008).

Sequential exposure to opposing force fields or visuomotor transformations in blocked schedules often leads to anterograde and retrograde interference (interference is anterograde when the preceding task interferes with the subsequent task, while it is retrograde when the subsequent task interferes with the memory of the preceding task). These interferences create large motor errors whenever the block alters (Brashers-Krug et al., 1996; Caithness et al., 2004; Gandolfo et al., 1996; Karniel & Mussa-Ivaldi, 2002; Krakauer et al., 2005; Wigmore et al., 2002), although re-adaptation is often faster with repetitive blocks (a phenomenon called savings) (Heald et al., 2021; Herzfeld et al., 2014; Oh & Schweighofer, 2019; Sugiyama et al., 2023; Turnham et al., 2012). Because of such interference, it has been claimed that dual adaptation is impossible (Gupta & Ashe, 2007; Hinder et al., 2008).

However, several studies have shown that humans are capable of overcoming interference by consolidating the motor memories of multiple environments and immediately switching among them by randomly presenting the multiple environments and/or providing additional environmental contexts (Forano & Franklin, 2020; Forano et al., 2021; Heald et al., 2021; Hinder et al., 2008; Howard et al., 2013; Krouchev & Kalaska, 2003; Lee & Schweighofer, 2009; Magnard et al., 2024; Osu et al., 2004; Shelhamer et al., 2005; Wada et al., 2003). Consolidation here is defined as resistance to retrograde interference (Caithness et al., 2004; Krakauer et al., 2005), assuming that motor memory is transformed from a fragile to a more stable state (Albouy et al., 2013; Thurer et al., 2018), and switching as the effective retrieval of the saved motor memory corresponding to presented contextual cues, being susceptible neither to anterograde nor retrograde interference (Zarahn et al., 2008). In particular, simultaneous learning of two opposing environments is possible when these conflicting environments are presented randomly with contextual cues (Forano & Franklin, 2020; Forano et al., 2021; Lee & Schweighofer, 2009; Osu et al., 2004; Shelhamer et al., 2005; Wada et al., 2003), although simultaneous adaptation is slower than adaptation for each task. Given these previous reports of both the ability and the inability to learn, we propose that there must be memory mechanisms that can adequately explain these contradictory results.

When adapting to novel environments, motor memories that capture the relationships between the desired behavioral consequences and the motor commands are formed as internal models in the central nervous system (CNS) (Kawato, 1999; Shadmehr & Mussa-Ivaldi, 1994; Shadmehr et al., 2010; Wolpert et al., 1998). The internal models are consecutively updated based on errors in preceding trials (Franklin et al., 2008; Herzfeld et al., 2014; Lee et al., 2018; Mattar & Ostry, 2007; Oh & Schweighofer, 2019; Scheidt et al., 2001; Takahashi et al., 2001). Smith et al. (2006) proposed a computational model that comprises a fast-learning, fast-forgetting memory process and a slow-learning, slow-forgetting memory process; see also (Coltman et al., 2019; Huberdeau et al., 2015; McDougle et al., 2015; Sing & Smith, 2010; Turnham et al., 2012). This fast/slow model accounted for several experimental phenomena in motor adaptation, including anterograde interference, spontaneous recovery, and rapid unlearning. However, this model cannot by itself reproduce the simultaneous acquisition and switching of multiple motor memories because a motor memory corresponding to each environment must be acquired to learn multiple environments simultaneously (Forano & Franklin, 2020; Haruno et al., 2001; Heald et al., 2021; Lee & Schweighofer, 2009; Oh & Schweighofer, 2019; Wolpert & Kawato, 1998). The MOSAIC (Modular Selection and Identification for Control) model was proposed to explain adaptation to multiple environments (Haruno et al., 2001; Wolpert & Kawato, 1998). Lee and Schweighofer (Lee & Schweighofer, 2009) then proposed a model with a single fast-learning, fast-forgetting “generalist” process and multiple slow-learning, slow-forgetting “specialist” processes, which were protected from interference.

In MOSAIC and recent extensions, selection between multiple models and update of each model depends on responsibility signals that combine three factors (Haruno et al., 2001; Heald et al., 2021): the prior history of the perturbation, possible sensory cues, and following feedback, the likelihood of the model, which depends on the sensory prediction error for each model weighted by the spatial precision of each model (i.e., the inverse of its width)(Oh & Schweighofer, 2019). Such models learn multiple environments simultaneously when information such as context or prediction error is provided and thus cannot sufficiently explain the observed behaviors of greater interference in the block than in random schedules. In addition, these models did not account for the large individual differences in learning in multiple environments.

In recent work (Oh & Schweighofer, 2019), we showed that interindividual differences in the rate of de-adaption and re-adaption to a visuomotor rotation depended on the ability to create and update new internal models specific to the perturbations and then easily switch between models (resulting in fast de-adaptation and re-adaptation), or to continuously update an existing model (resulting in slower de-adaptation and re-adaptation). These interindividual differences were controlled by the relative precision of the different models, which yielded individual differences in model selection and learning rates by modulating the responsibility signals.

Here, we hypothesize that individual differences in both the skill level and rate of skill acquisition in dual-adaptation paradigms largely arise from the ability to learn and switch between multiple tasks simultaneously. Combining MOSAIC and our previous single generalist and multiple specialists model (Lee & Schweighofer, 2009), we propose a new model equipped with two types of memory architecture: one with a single generalized memory store, which assumes continuity of the environment over trials as in block schedules, and the other with multiple memory stores specific to each environment, which assume that the environment may change between trials, as in random schedules. We further hypothesized that individual differences in model precisions (i.e., widths) and learning rates between the generalist memory module and the specialist modules can explain both population-wise and individual-wise characteristics of motor learning.

We first performed a series of multi-day force-field dual-adaptation experiments with block or random schedules. We examined the ability to learn and retain both tasks in 1-day retention tests, as well as individual differences in learning and retention, following these two practice schedules. We then simulated the different training schedules to examine whether the proposed model successfully accounted for poor memory consolidation and inappropriate switching after block presentation and superior memory consolidation and successful switching after random presentation, as well as large individual differences in random schedules.

## Results

Participants learned reaching movements to eight targets located radially from a central start position. The movements occurred in either a clockwise (CW) or counterclockwise (CCW) velocity-dependent rotational force field (Figure 1). After the presentation of audiovisual cues that indicates the direction of rotation, one of the eight targets was randomly presented. Participants were required to reach the target in a straight trajectory. Participants learned two tasks, CW and CWW, in either block schedule or random schedule. Participants who practiced in a block schedule for four consecutive days were tested in either a block schedule (BLOCK-BLOCK group) or a random schedule (BLOCK-RANDOM group) after the last block training (Table 1).

**Figure 1:**
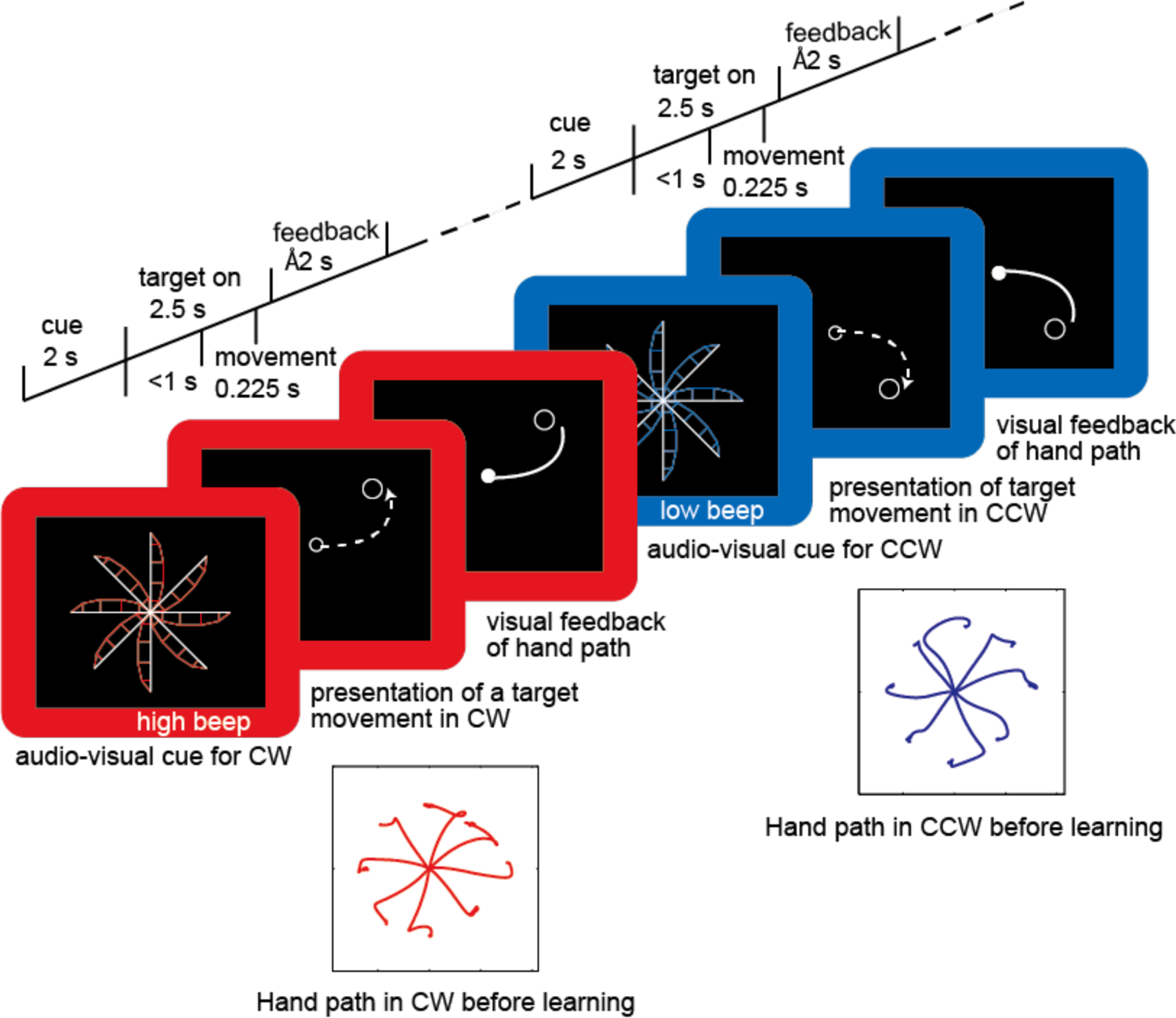
Experimental procedure, with an example of two trials in the random condition. Each trial consisted of presenting audio-visual cues, target presentation, movement, and visual feedback of the movement trajectory. In RANDOM training and test sessions, the order of presentation of the two force fields was random. The insets show examples of hand paths at the initial exposure to CW (red) and CCW (blue) force fields, respectively.

**Table 1.** Experimental protocol. The numbers in parenthesis indicate the number of trials in each session. In block training and test sessions, half of the participants were exposed to blocks in the reverse order except for training sessions of the BLOCK-RANDOM group in which all participants were exposed in the order of CW-CCW-CW-CCW.

|  |  |
| --- | --- |
| BLOCK-BLOCK group (8 participants) |  |
| Train | Day 1: CW (112 trials)<br>Day 2: CCW (112 trials)<br>Day 3: CW (112 trials)<br>Day 4: CCW (112 trials) |
| Test | Day 4: CCW (64 trials) – CW (64 trials) – CCW (64 trials) |
| BLOCK-RANDOM group (11 participants) |  |
| Train | Day 1: CW (112 trials)<br>Day 2: CCW (112 trials)<br>Day 3: CW (112 trials)<br>Day 4: CCW (112 trials) |
| Test | Day 4: Random (224 trials) |
| RANDOM-RANDOM group (10 participants) |  |
| Train | Day 1: Random (224 trials)<br>Day 2: Random (224 trials) |
| Test | Day 2: Random (256 trials with 32 catch) |
| RANDOM-BLOCK group (12 participants) |  |
| Train | Day 1: Random (224 trials)<br>Day 2: Random (224 trials) |
| Test | Day 2: CCW (64 trials) – CW (64 trials) – CCW (64 trials) |
| Control Experiment |  |
| BLOCK control group (10 participants) |  |
| Baseline | CCW (64) – CW (64) – CCW (64) |

Participants who practiced in a random schedule for two consecutive days were tested in either a block (RANDOM-BLOCK group) or a random schedule (RANDOM-RANDOM group). During RANDOM training sessions, the number of trials for each force field (112) was the same as the number of trials in BLOCK training sessions. All groups executed 448 trials in total during training, with 224 CW trials and 224 CCW trials. A control group was presented with only a test block session (BLOCK-control group). The task and feedback in the test blocks were the same as those in the training blocks. Adaptation was assessed by deviations from the straight path computed as the signed areas between the actual hand path and the line joining the start and target centers, with a positive sign indicating a CCW hand path deviation and a negative sign indicating a CW deviation (directional error). These errors were averaged for each cycle, whereby one cycle consisted of eight consecutive trials in one of the two force fields, including movements to all eight targets.

### Block training induced interference

We first investigated the effect of block schedule training on performance in the following block or random test session. When participants were initially exposed to the CW force fields, hand trajectories were highly distorted and curved in the direction of the applied force (Figure 1, inset). The movement error measured by directional error (see Methods) was reduced in an exponential-like manner as practice proceeded (Figure 2a, b). When exposed to an opposite CCW force field on Day 2, participants produced a larger magnitude of directional error than on Day 1, as in previous studies (Caithness et al., 2004; Krakauer et al., 2005). On Days 3 and 4, the magnitudes of initial errors were as large as or larger than those on Day 1. The magnitude of the directional error approached zero in an exponential-like manner at the end of each daily training block.

**Figure 2:**
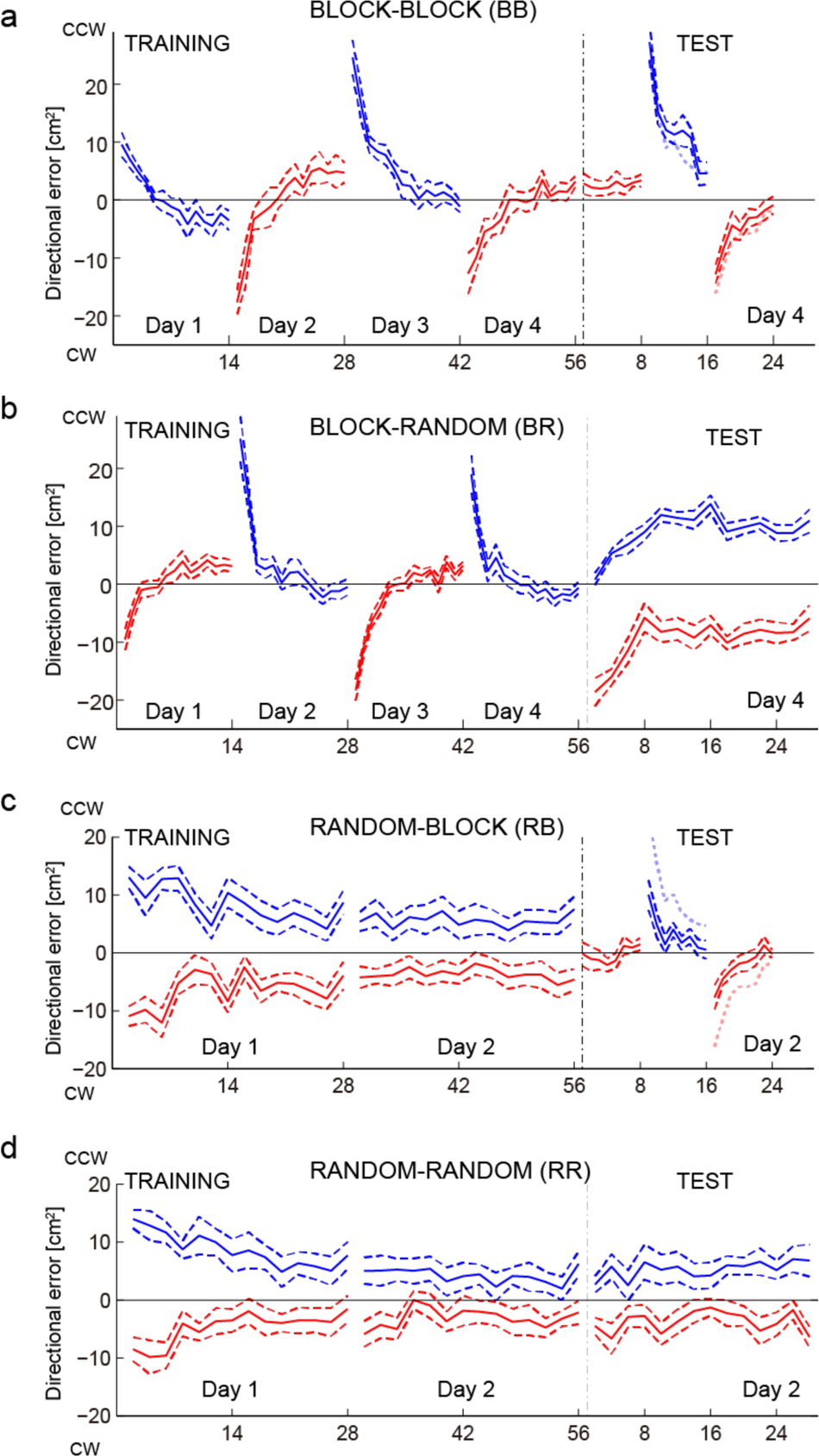
Learning curves for the BLOCK-BLOCK (a), BLOCK-RANDOM (b) RANDOM-BLOCK (c), and RANDOM-RANDOM (d) groups. Average directional errors and s.e.m (dashed lines) in CW (red) and CCW (blue) across participants are plotted against cycles of 8 targets for TRAINING and TEST. The sign was flipped for the participants who were exposed to blocks in reverse order before averaging. The thick dotted curves with a light color in TEST denote average directional errors of the second (blue) and third (red) blocks of the BLOCK-control group.

The observed large aftereffects and considerable re-adaptation process demonstrate anterograde and retrograde interference, which are incompatible with the possibility of the consolidation and switching of motor memory learned on Day 1 and 2.

In the following block test session on Day 4 in the BLOCK-BLOCK group, the magnitude of the directional error was small in the first block (Blk 1 in Figure 2a) in the test session because the direction of the force field was the same (CW) as that in the last training session block. However, the error increased when the force field switched to the CCW block (Blk 2 Figure 2a). The error in the subsequent CW block (Blk 3 in Figure 2a) increased again. The learning curves in the second and third test session blocks were comparable with those in the BLOCK-control group in which participants only experienced the block test session (pale blue and pale red dotted curves), suggesting that the preceding block training session was not effective for memory consolidation or switching of multiple environmental dynamics.

In the following random test session in the BLOCK-RANDOM group, the magnitude of the initial directional error was small when CCW was presented and large when CW was presented (Figure 2b). Good initial performance in CCW test session trials reflected memory preservation from the preceding CCW training block. Poor initial performance in CW test session trials suggested that the memory of the CW training block presented the day before was not preserved. Consequently, although participants had already experienced both force fields in blocks, the initial directional error for CW in the random test session was worse than for the first exposure to CW force field in either block or random schedules (Figure 2c and d). As the test proceeded, performance in CCW worsened in parallel with the gradual improvement in performance in CW, reflecting probable learning of the average of the two dynamics rather than learning them separately, as previously reported (Scheidt et al., 2001).

### Random training reduced interference

We next investigated the effect of training in random presentation on performance in the following block (Figure 2c) or random (Figure 2d) test session. In both RANDOM-BLOCK and RANDOM-RANDOM groups, the average performance was superior in test sessions compared to that in BLOCK-BLOCK and BLOCK-RANDOM groups, respectively (compare Figure 2c to Figure 2a, and Figure 2d to 2b).

To summarize performance, we defined the difference errors as the difference in the directional errors between the CW and CCW (CCW - CW), where larger positive values indicated poorer adaptation, and negative values indicated over-adaptation. Figure 3a compares the magnitude of difference error computed from the second and third blocks in the test session of BLOCK-BLOCK, RANDOM-BLOCK, and BLOCK-control groups. The error was significantly smaller only in the RANDOM-BLOCK group (Kruskal-Wallis test, p < 0.001; post-hoc Wilcoxon test, p < 0.01, effect size r > 0.6), suggesting that block training did not result in consolidation and switching of motor memory responsible for each force field. Results also confirmed the preservation of multiple motor memories in the block presentation after random training. The difference error across cycles in random test sessions was significantly smaller in the RANDOM-RANDOM group than in the BLOCK-RANDOM group (Wilcoxon rank sum test, p = 0.018, effect size r > 0.5; Figure 3b), suggesting that memory consolidation and effective switching occurred after random training but not after block training. Note that although RANDOM-RANDOM group received only two days of training, performance was still much better in test than BLOCK-RANDOM group who received four days for training.

**Figure 3:**
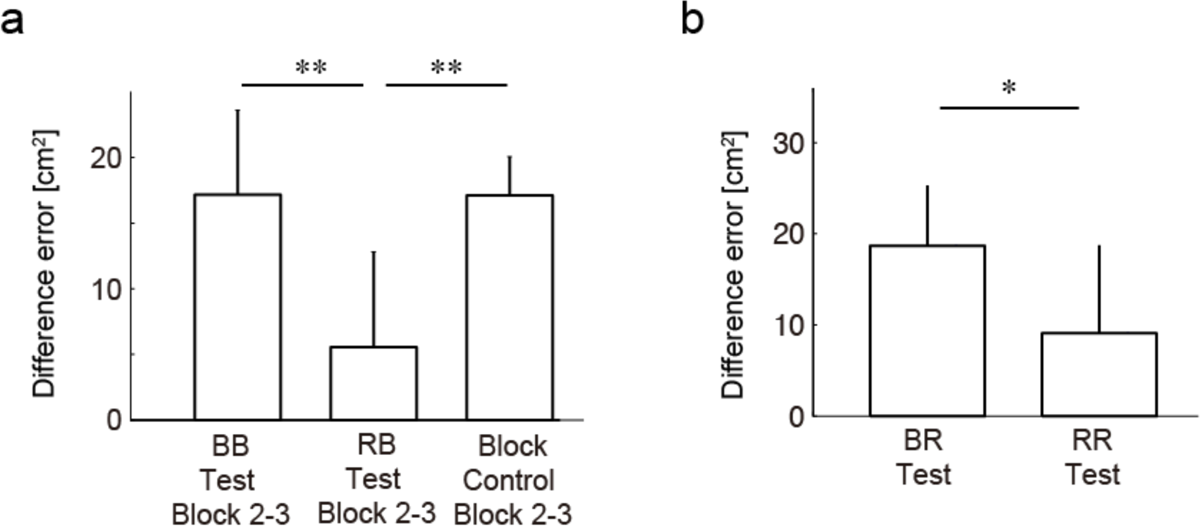
(a) Comparison of average difference errors across participants and s.e.m from the second and third blocks during the test session among BLOCK-BLOCK (BB), RANDOM-BLOCK (RB), and BLOCK control groups. Errors in the RB test session were significantly smaller than in the other two groups (** denotes p < 0.01). (b) Comparison of average difference errors during the test session and s.e.m between BLOCK-RANDOM (BR) and RANDOM-RANDOM (RR) groups. Errors in the RR test sessions were significantly smaller than those in the BR test session (* denotes p < 0.05).

### Individual differences in random learning

To examine interindividual variability in performance, we compared the distribution of difference errors between test sessions after random training and those after block training (Figure 4a). Difference errors were distributed with significantly more dispersion after random training than after block training (Ansari-Bradley one-tailed test of equal variance, p = 0.039). Coefficient of variations (standard deviation divided by the mean) of the difference errors and their bootstrap confidence intervals (95% CI) after random and block training were 86.8% (58.5, 153.1) and 33.5% (24.0, 50.4), respectively, showing larger dispersion after random training. These results and those of Figure 3 show that, overall, performance was poorer after block training than after random training but also more variable after random training.

**Figure 4:**
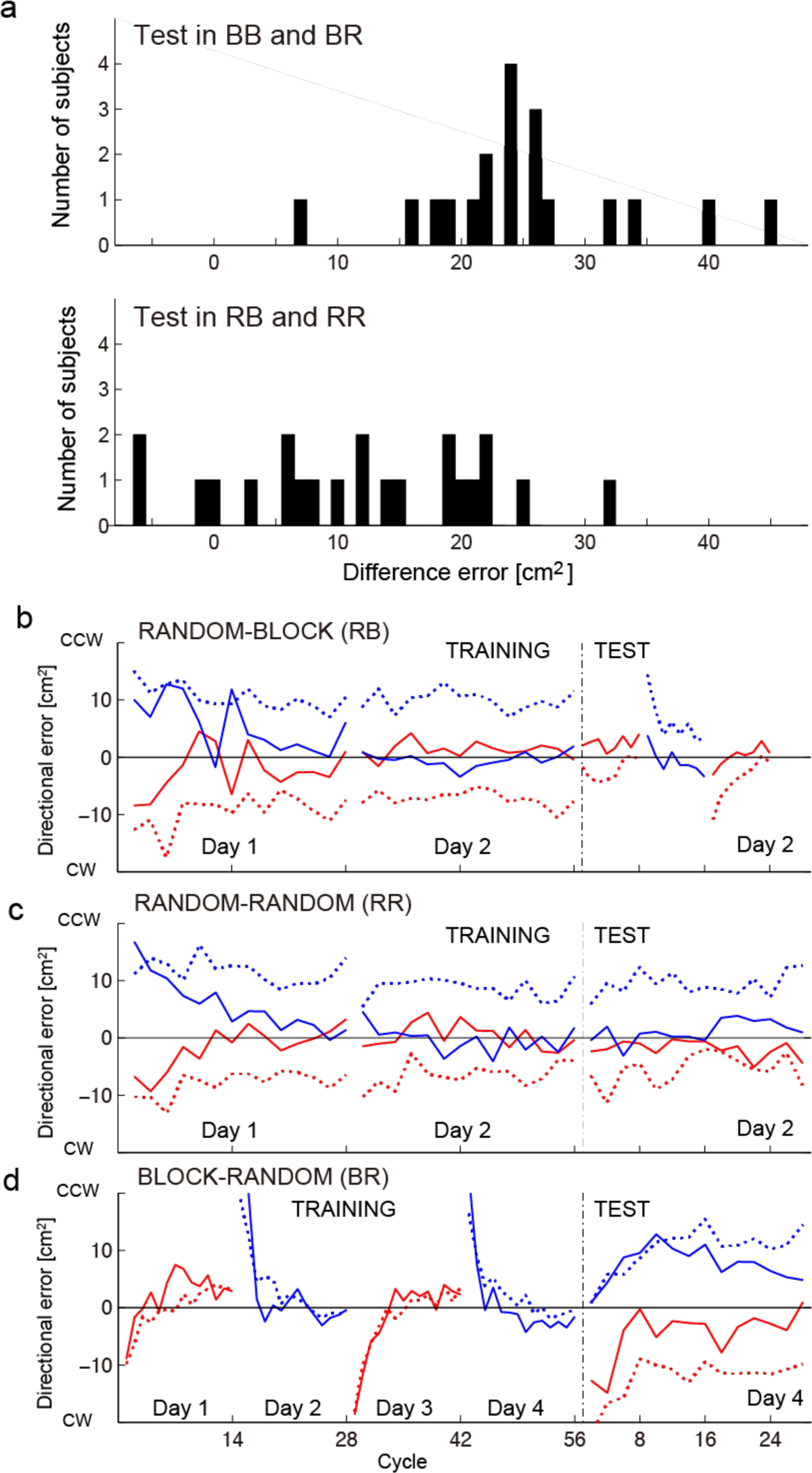
Individual differences in learning under random presentation. (a) Histogram of the difference errors in test sessions of BLOCK-BLOCK and BLOCK-RANDOM groups (upper panel), and RANDOM-BLOCK and RANDOM-RANDOM groups (lower panel). Negative difference errors indicate over-adaptation. (b-d) Learning curves of good learners (solid curves) and poor learners (dotted curves) separately presented for RANDOM-BLOCK (RB), RANDOM-RANDOM (RR), and BLOCK-RANDOM (BR) groups. For RB and RR groups, participants were classified based on difference errors in the second half of the training session by the threshold of the mean (8.42 cm^2^). Five out of 12 participants in the RB group and five out of 10 participants in the RR group were denoted as good learners. For the BR group, participants were classified based on difference errors in the last two cycles (13 and 14 cycles) by the threshold of the mean (17.00 cm^2^). Four out of 11 participants were denoted as good learners.

We separated the 22 participants assigned to random presentation training (RANDOM-BLOCK and RANDOM-RANDOM groups) into good or poor learners based on the mean difference error during the training session on Day 2 (cycles 29 to 56) (Figure 4b and c). Because their mean difference error on Day 2 training was smaller than the average of 22 participants, we classified ten participants (five per group) as good learners (solid blue and red lines). The other 12 participants were assigned as poor learners (seven for RANDOM-BLOCK, five for RANDOM-RANDOM; dotted blue and red lines). As shown in Figure 4b and c, participants who learned well in the random training session performed well in the subsequent block or random test session, while those who learned poorly in the random training session performed poorly in the subsequent block or random test session (Wilcoxon rank sum test, p < 0.01). There was a significant correlation between difference errors on Day 2 random training and test sessions for both RANDOM-BLOCK (Spearman ρ = 0.84, p < 0.001) and RANDOM-RANDOM (Spearman ρ = 0.85, p < 0.01) groups.

We then separated the 11 participants in the BLOCK-RANDOM group into good learners (four participants) and poor learners (seven participants) based on the mean difference error during the last four cycles (25 to 28) of random test sessions (Figure 4d). The mean difference error of the preceding block training session on Day 3 and 4 (cycles 29 to 56) was not correlated with the final performance in the last four cycles of the random test session (Spearman ρ = 0.23, p = 0.50; n.s.), indicating that the memory shaped during subsequent random training was independent of the memory shaped during the preceding block training.

### The ratio of learning rates of the generalist to specialist memory modules explained behavioral results: a simulation

To account for the experimental results, we performed simulations based on the MOSAIC architecture, as it can learn and switch among multiple internal models (Wolpert & Kawato, 1998). In MOSAIC, each module *i* consists of a pair of forward and inverse models and is selected and updated based on the responsibility signal *λ_i_* (*t*). In simulations, we defined a simplified environment of the task under CCW and CW force fields. The position of the cursor and force applied at the cursor were represented by [*x*(*t*) *y*(*t*)]^T^ and [*u_x_* (*t*) *u_y_* (*t*)]^T^, respectively. The equation of motion was defined by:

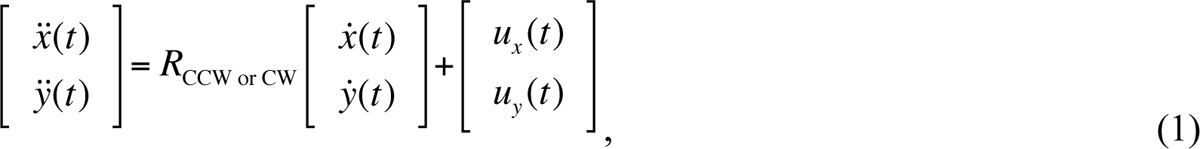

where *R*_CCW or CW_ are the rotation matrices; 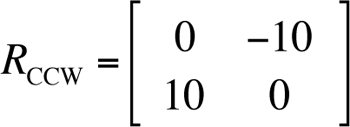 and 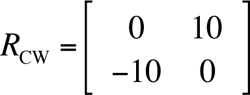. The mass of the arm tip was 1 kg for simplicity. The state variable and motor output were defined as 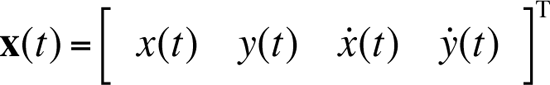 and 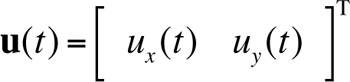, respectively. The following linear forward models predicted the state with each module *i*:

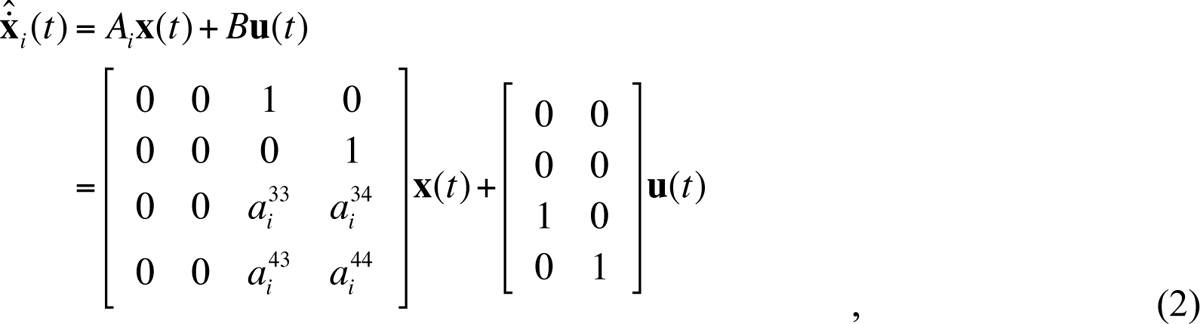

where *A_i_* and *B* are the forward model parameter of module *i*. The motor output of each module was computed by inverting the forward models,

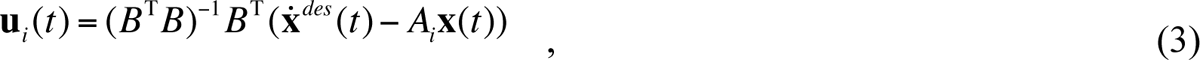

where the desired trajectory **x** *^des^* (*t*) was set as minimum jerk trajectory. The final motor output **u**(*t*) is the summation of the control signal from each module **u** (*t*) weighted by its responsibility signal *λ_i_* λ(*t*).

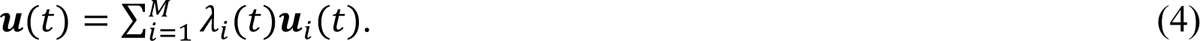

In parallel, the internal forward model parameters in *A* of each module *i* are updated according to the gradient method.

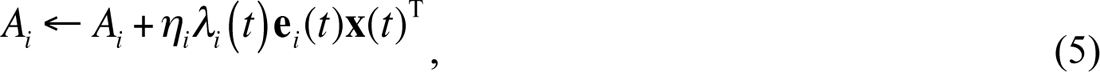

where *η_i_* is the learning rate of the module *i*. Note that the effective learning rate for each module is given by *η_i_λ_i_* (*t*).

The responsibility signal *λ_i_* (*t*) is computed from prediction error **e***_i_* (*t*) between the actual state (velocity) **x**^&^ (*t*) and the state predicted by the forward model X^^^_i_(*t*), and prior probability of the responsibility signal λ*_i_* (*t*), as follows:

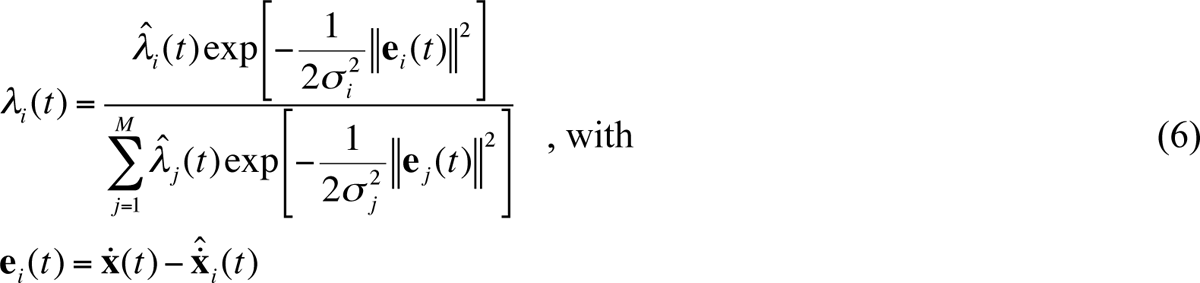

where the parameter *σ* _i_^2^ determines the selectivity spatial width of switching in responsibility signal of module *i*, and *M* is the number of modules. The prior probability for each module is given by:

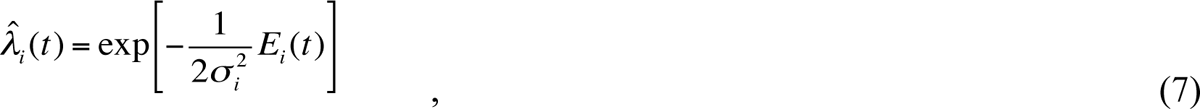

with *E_i_* (*t*) is a temporally smoothed prediction error. The characteristics of the prior probability of the responsibility signals λ*_i_* (*t*), and therefore that of *E_i_* (*t*), is a key factor for learning in multiple environments (Imamizu et al., 2007). The temporal continuity of the environment is informative in recommending not to frequently switch, and to stabilize network performance. However, in random presentation of the opposing force fields, environments are discontinuous across trials and continuous within a trial. Thus, the generalist module is expected to deal with a single environment that will continue to exist for at least a block of trials. In contrast, the specialist modules are expected to collectively deal with multiple environments that appear stochastically at each trial. Therefore, in the present model, in addition to a generalist module with *E* (*t*) that exhibits temporal continuity across trials (like the modules in MOSAIC), we assumed the existence of specialist modules whose *E* (*t*) were smooth within a trial but reset on the next trial (Equation 8). The temporally smoothed prediction errors of module *i* at time *t* are therefore described as

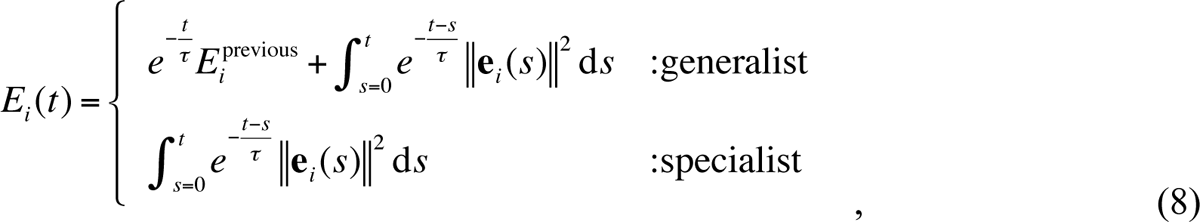

where the parameter *τ* is a time constant for temporal smoothness and was fixed to 1 s for both generalist and specialist modules in the present simulation, **e***_i_*(*s*) is the prediction error at time *s* (Equation 6), and *E* ^previous^ denotes *E* at the end of the previous trial.

In our simulation of dual adaptation, we prepared a single generalist module and two specialist modules with forward model parameters of A*_generalist_*, A*_specialist_*, and A*_specialist_* similarly to (Lee & Schweighofer, 2009). Each module computes a control signal based on Equation 3. We randomly searched for the learning rate *η_i_* in Equation 5 and the width of switching in the responsibility signal *σ* ^2^ in Equations 6 and 7, for each type of module so that the model could reproduce the experimentally observed characteristics of good learners.

Two requirements needed to be met for the reproduction of good learners. First, the model needed to account for the retention of motor memories corresponding to two environmental dynamics and effective switching in test sessions after random training in RANDOM-BLOCK and RANDOM-RANDOM conditions. Second, the model needed to account for poor retention or switching in test sessions after block training in BLOCK-BLOCK and BLOCK-RANDOM conditions. We defined the threshold of simulated maximum magnitude of directional error (error magnitude) as less than 10 cm^2^ for successful retention. After random training, the error magnitude of test sessions had to be less than the threshold to replicate good memory retention in good leaners; in contrast, after block training, the error magnitude of test sessions had to be large and above threshold to replicate poor memory retention, even in good learners. Assuming that the parameters of two specialists were the same, we randomly generated 50,000 of four parameter sets consisting of *η_generalist_*, *η_specialist_ σ* ^2^*_generalist_*, and *σ* ^2^*_specialist_*, simulated the four experiments (BLOCK-BLOCK, BLOCK-RANDOM, RANDOM-BLOCK, and RANDOM-RANDOM) using these parameter sets.

Of 50,000 parameter sets, 452 satisfied the two above conditions and thus reproduced the behavioral characteristics of good learners. The learning rate *η_i_* was significantly larger for the generalist module than for the specialist module (Wilcoxon signed rank test, p < 0.0001; see histograms in Figure 5a and b). That is, the generalist module exhibited faster learning, whereas the specialist module exhibited slower learning. Thus, in our simulations, fast and slow learning processes emerged spontaneously even though we did not explicitly preset fast and slow processes beforehand (Sing & Smith, 2010; Smith et al., 2006). Similarly, the width parameter*σ* ^2^ was significantly larger for the generalist module than for the specialist module (Wilcoxon signed rank test, p < 0.0001, see histograms in Figure 5c and d). This indicated that the generalist module was selected even when its prediction error was large, whereas each specialist module was selected when its prediction was accurate.

**Figure 5:**
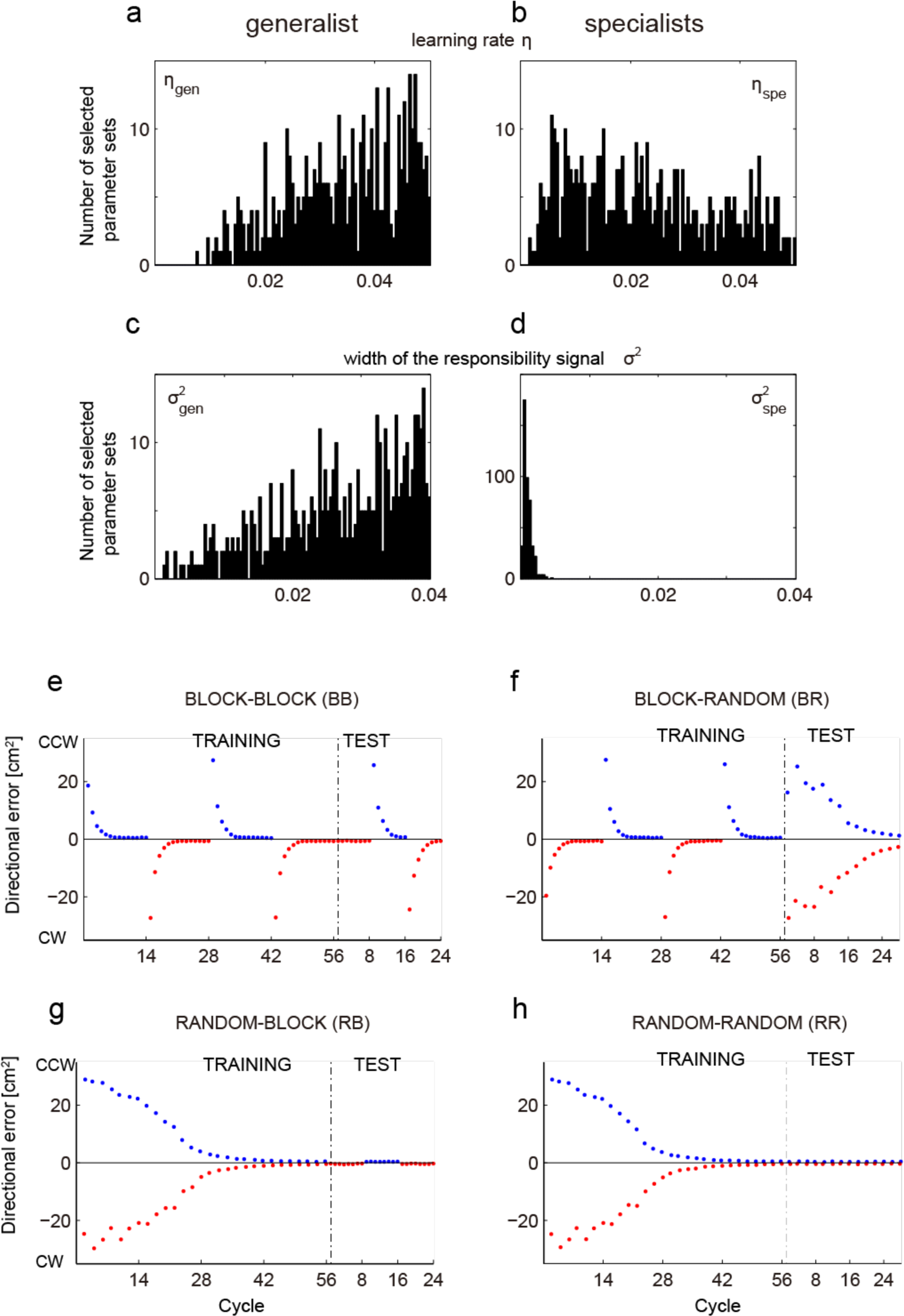
Parameter sets of the modified MOSAIC model with one generalist module and two specialist modules that explained human behavior in our experiment, i.e., no consolidation in BB and BR test sessions and good consolidation in RB and RR test sessions. (a-d) Histograms of learning rate *η* for generalist (a) and specialist modules (b), and histograms of width of switching in responsibility signal *σ* ^2^ for generalist (c) and specialist modules (d) from selected parameter sets. (e-h) Examples of simulated learning curves using one parameter set that satisfied human behavior. Directional error was plotted against cycles in the same way as the human data shown in Figures 2 and 3 for BB (e), BR (f), RB (g), and RR (h) conditions.

Considering the effective learning rate *η_i_ λ_i_* _(_*t*_)_ (Equation 5), the sharp switching (smaller *σ* ^2^) in specialists resulted in even slower learning of specialists, since a specialist easily leaves out of the selection window for the responsibility signal. This corresponds to the observed slow learning in random presentations and is consistent with the results from a previous model with multiple slow states (Lee & Schweighofer, 2009) that learned each environment separately. Similarly, large *η^2^_i_* and *σ* in the generalist module are consistent with the fast memory state in this previous model.

An example of the simulated learning curves using one of the parameter sets that reproduced all four behavioral results of BB, BR, RB, and RR is depicted in Figure 5 (lower panels). The simulation qualitatively reproduced the time course of errors of the good learners under all four conditions.

Figure 6 plots the evolution of the responsibility signal *λ_i_* (*t*) using the same parameter set as that in the example of Figure 5. In the BLOCK-BLOCK condition, the responsibility signal of the generalist rapidly increased in each block (magenta curves in Figure 6a). In the BLOCK-RANDOM condition, the responsibility signal of specialists started to evolve when random test sessions were introduced and then gradually overwhelmed the generalist module that was dominant in the block training session (Figure 6b). In the RANDOM-BLOCK and RANDOM-RANDOM conditions, the responsibility signal of the generalist module gradually decreased, while that of the specialist modules evolved during training sessions and remained high in the test period (cyan curves in Figure 6c and d).

**Figure 6:**
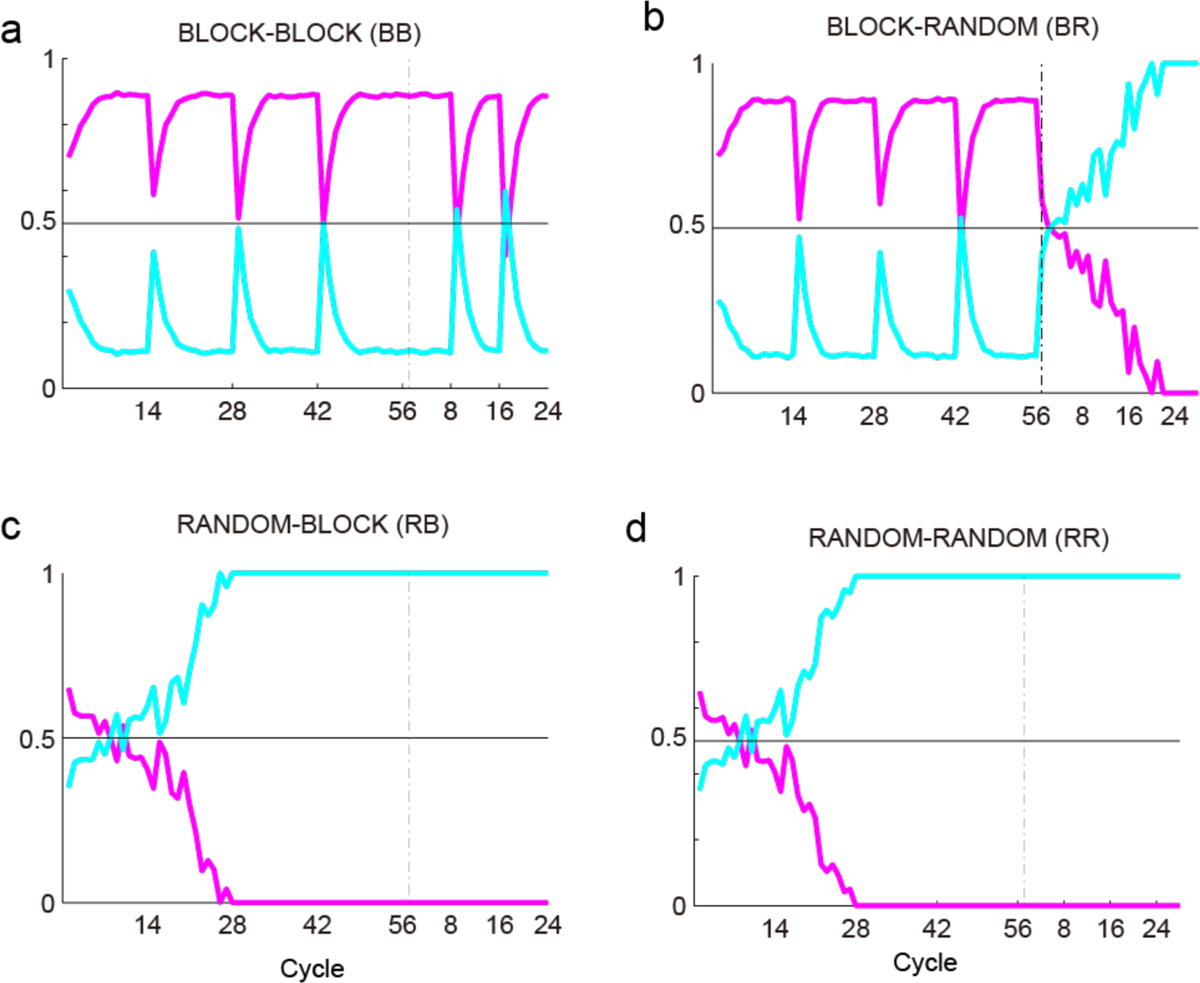
Evolution of responsibility signals using the parameter set of Figure 5. Magenta curves denote the responsibility signal of the generalist. Cyan curves denote the sum of the responsibility signals of the two specialists. When the magenta curve reaches 1 (a,b), only the generalist module is selected to control the hand. When the cyan curve reaches 1 (c,d), one of the two specialist modules is selected. When neither reaches 1, the hand is controlled by the weighted summation of generalist and specialist modules.

To examine the relationship between the characteristics of the parameter sets and consolidation, we plotted the maximum magnitude of directional error (error magnitude) in test sessions against the ratio of the effective learning rate between generalist and specialist modules. It can be shown (see Methods) that the effective learning rates are proportional to *η_generalist_*, *η_specialist_ σ* ^2^*_generalist_*, i.e., the ratio of learning rate *η* between generalist and specialist modules, multiplied by the ratio of width of responsibility signal *σ* ^2^ between generalist and specialist modules. The boxplots in Figures 7a and b show the distribution of error magnitude for all 50,000 parameter sets in the simulation of BLOCK-BLOCK, RANDOM-BLOCK, BLOCK-RANDOM, and RANDOM-RANDOM for different such ratios.

**Figure 7:**
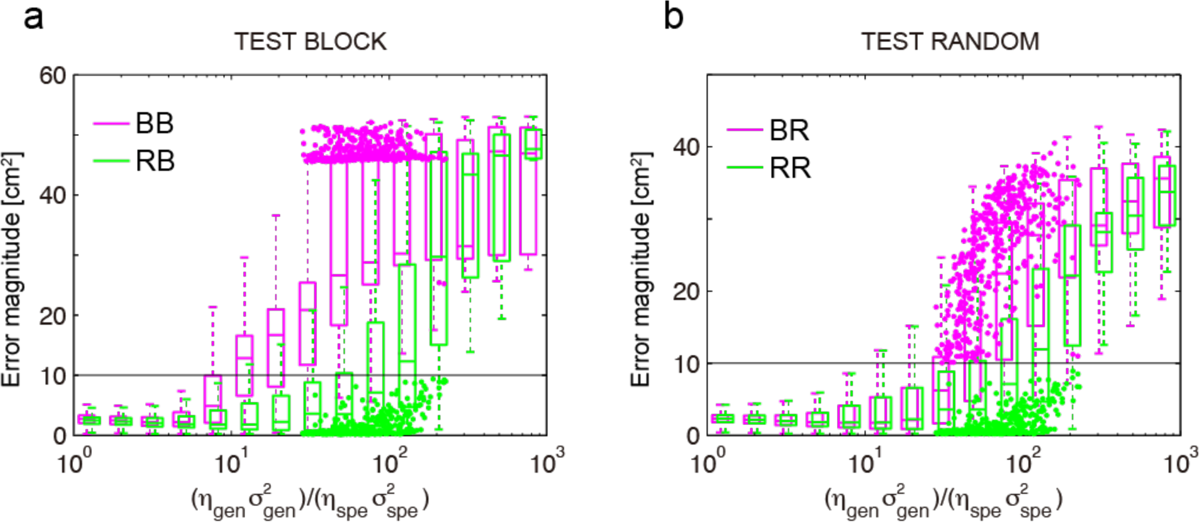
Maximum magnitude of directional error (error magnitude) in test sessions plotted against the ratio of effective learning rates of generalist to specialist (*η_generalist_*, *η_specialist_ σ* ^2^*_generalist_*), in simulations of TEST BLOCK (a) and TEST RANDOM (b) following both training conditions RANDOM and BLOCK. The black horizontal line at 10 cm^2^ indicates the threshold of error magnitude for successful consolidation in the tests. The boxplots show the median, 25^th^, and 75th percentiles of error magnitude for all 50,000 parameters for 10^0.2^ bins, for increasing values of the effective learning rate ratio. The dots show the error magnitude for the good learners corresponding to the 452 selected parameter sets. The colors correspond to the type of training, with magenta for BLOCK and green for RANDOM. Thus, in a), magenta indicates parameters for BLOCK-BLOCK (BB) and in b) for BLOCK-RANDOM (BR). In a) green indicates parameters for RANDOM-BLOCK (RB) and in b) for RANDOM-RANDOM (BR). When the ratio of effective learning rates was less than ∼20, retention occurred following both training conditions (as the test error magnitude is smaller than the threshold in RB and RR), contradicting the data. A ratio of ∼50 to ∼130 corresponds to good learners as shown by the distribution as the median of the test error magnitude is below the threshold following random training and above the threshold following block training. Larger ratios correspond to poor learners, as the test error magnitude is greater than the threshold in RB and RR.

When the generalist learned with similar or slightly faster speed than that of specialists (approximately) 1 to 20 times faster, (i.e., when the ratio of effective learning rate was 10^0^ < (*η_generalist_*, *η_specialist_ σ* ^2^*_generalist_*), < 2 x10^1^ on the abscissa), the error magnitude of all conditions was smaller than the threshold of 10 cm^2^, indicating retention in all four conditions, which contradicted all observed behavioral results.

Larger ratios accounted for the data. As indicated by the dots showing the error magnitude for each of the selected 452 parameter sets in Figure 7a and b, the behavior of good learners was accounted for when the generalist learned 50 to 130 times (approximately) faster than the specialists (5 x 10^1^ < (*η_generalist_*, *η_specialist_ σ* ^2^*_generalist_*), < 13 x 10^1^ on the abscissa). In this case, the median of the error magnitude and that of the selected parameters in BLOCK-BLOCK and BLOCK-RANDOM conditions increased (magenta plots in Figures 7a and b: no consolidation), while that of RANDOM-BLOCK and RANDOM-RANDOM conditions stayed below the threshold (green plots in Figures 7a and b: consolidation). The behavior of poor learners corresponded occurred when the generalist learned (approximately) more than 200 times faster than the specialists (2 x 10^2^ < (*η_generalist_*, *η_specialist_ σ* ^2^*_generalist_*), on the abscissa). In this case, the median of the error was higher than the threshold in any condition, corresponding to the behavior of poor learners.

In addition, as observed in the behavior (Figure 4a), the errors were distributed with significantly more dispersion after random training (RANDOM-RANDOM and RANOM-BLOCK) than after block training (BLOCK-BLOCK and BLOCK-RANDOM) in the parameter range that corresponds to both good and poor learners (i.e., the generalist learned 50 to 800 times faster) (Ansari-Bradley one-tailed test of equal variance, p < 0.001). The appropriate ratio of the effective learning rate successfully explained the average performance difference between block and random training, and its distribution explained individual performance differences in random training.

## Discussion

Our results show that, as previously reported, all participants in the block training schedule adapted to the force field presented in a block but showed large interference in the subsequent opposing force field blocks; thus, adapting to the two force fields was impossible. In contrast, participants in the random training schedule could, as a group, adapt to the two conflicting tasks simultaneously. This indicates that the motor memories of the two perturbations were not consolidated separately after block presentation. In contrast, the motor memories were consolidated after random presentation. Such consolidation improved the switching performance among the two environments in subsequent test sessions with both block and random presentations. In addition, while we observed little variability in learning performance and retention in block training across participants, we observed large inter-individual differences in random training: some participants showed almost perfect learning of both tasks, whereas others showed minimal learning.

A modified MOSAIC model equipped with one generalist module and two specialist modules was able to account for both the different behaviors in the two conditions and the large variability in random schedules for adequate parameter ranges. Crucially, for the selected parameters, the width (inverse of precision) of switching in responsibility signals was larger in generalist than in specialist modules. That is, a generalist module was selected even when errors were large and by learning quickly, attempted to cover any environment. Conversely, a specialist module was selected only when the module accurately predicted the environment and learned slowly to be a specialist in a given environment. In addition, at the group level, the greater effective learning rates assigned to the generalist than to the specialists agreed qualitatively with the difference in learning rates of previously proposed fast memory and slow memory processes (Lee & Schweighofer, 2009; Smith et al., 2006). Instead of including a single generalist, Forano and Franklin (2020) proposed a model with multiple parallel motor memories, each involving a fast, slow, and ultraslow process, all weighted by a responsibility estimator. They showed that their model better explains the spontaneous recovery after dual adaptation in a block schedule. Whereas their model could explain the individual difference by variability in learning rates, we believe that it could not account for our results in block schedules because of the lack of a generalist module.

Here, we assumed that the prior distribution of the responsibility signal for specialists was reset for each trial, unlike the prior distribution for the generalist. This discontinuity in the specialist facilitated the selection of specialists in discontinuous environments, such as the random conditions, and was suppressed in continuous environments, such as the block conditions. Recent studies have reported that environmental consistency plays an essential role in determining the rate of motor adaptation (Gonzalez Castro et al., 2014; Herzfeld et al., 2014). It remains unclear whether resetting the prior responsibility distribution is a fundamental brain property and how this may be implemented.

An additional strength of the proposed model is its potential to explain individual differences. We observed smaller individual differences in block learning and larger individual differences in random learning (Figure 4); this contrast spontaneously emerged in simulations (Figure 7). When the ratio of effective learning rates between the generalist and specialist was intermediate, performance was poor in block training and good in random training as shown by good learners. When the ratio of effective learning rates between the generalist and specialist was large, performance was poor in both block and random training. Individual differences during skill acquisition have been examined in the field of applied psychology (Ackerman & Cianciolo, 2000) but have not been fully quantitatively investigated in computational motor control and learning research (but see (Ganesh et al., 2014; Magnard et al., 2024; Oh & Schweighofer, 2019; Takagi et al., 2017)). A consistent predictor or interindividual variability in motor learning is the capacity of spatial working memory (Anguera et al., 2010; Lingo VanGilder et al., 2018; Schweighofer et al., 2011), which may share neural resources with the fast-learning generalist module. In addition, individual differences have been examined in brain imaging studies (Bueti et al., 2012; Kanai & Rees, 2011), and interindividual variability in motor learning was reported to be correlated to differences in brain structures (Gelineau-Morel et al., 2012; Sampaio-Baptista et al., 2014; Vo et al., 2011). Future model-based fMRI studies are needed to relate our mechanistic explanations of good vs poor learners to such differences in neural activation or structures.

In our modified MOSAIC model, we did not introduce explicit contextual cue signals that directly controlled switching. Instead, we assumed that the magnitude of the errors that influence the responsibility signals would act as cues. We, however, are not implying that humans do not use contextual signals. For example, learning conflicting environments is facilitated when associated with synergetic cues such as posture and target location, which may facilitate the activation of specialist modules more easily than arbitrary cues (Forano et al., 2021; Gandolfo et al., 1996; Krakauer et al., 1999; Thomas & Bock, 2012; Tong et al., 2002; Woolley et al., 2007). Heald et al. (Heald et al., 2021) proposed a contextual inference model computing the probability that each known context, or a novel context, is active and creating a new memory when responsibility is high for the novel context. Our future framework should integrate such computation of contextual inference where probability will be influenced by the temporal aspect, i.e., how frequently the contextual signal changes.

Our results showing the benefit of a random practice schedule over block practice are consistent with previous studies in many types of motor tasks, including high-level cognitive tasks, and is termed contextual interference (CI) (Schmidt, 1988). Block conditions, which involve less CI enhance acquisition but degrade memory retention. Random training involving greater CI initially worsens performance and slows acquisition but promotes memory retention after a delay. In the model, the CI occurred because, in a block schedule, the generalist quickly improved performance, therefore reducing error-driven updates of the specialist processes. Because of interference in the generalist when another task was presented in the next block, poor long-term retention ensued. In random schedules, interferences in the fast process led to a slower change in performance, therefore increasing error-driven updates of the specialist and, thus, good long-term retention as in our proposed generalist and specialist architecture (see also, (Cross et al., 2007; Kim et al., 2015; Lage et al., 2015; Li & Wright, 2000; Schmidt, 1988; Schweighofer et al., 2011)).

A limitation of the current study was the exploratory nature of the analyses on individual variability; further hypothesis-driven experiments are required to confirm our results, although it is difficult to assign the participants a priori into good and poor learner groups. In addition, for modeling, the simulation environments were simplified and did not consider the nonlinear dynamics of the system. Furthermore, we prepared two specialists in advance of adaptation. A future model should not pre-specify the number of models, but the models should be created as needed (Heald et al., 2021; Oh & Schweighofer, 2019). The practical implication of our study is that individualized training programs can be provided according to the individual properties of memory systems when learning motor skills. Acknowledging the existence of individual differences in memory systems may be helpful for the practice of motor skill coaches or therapists.

## Methods

### Participants

In total, 51 healthy right-handed participants (18 to 38-years-old) participated in the study and were assigned to one of the five groups (Table 1). The institutional ethics committee of ATR approved the experiments. Participants provided written informed consent prior to experiments.

### Apparatus

Participants learned reaching movements to eight targets located radially from a central start position, as described previously (Osu et al., 2004). Movements occurred in either a clockwise (CW) or counterclockwise (CCW) velocity-dependent rotational force field produced by a manipulandum. The force fields are expressed as:

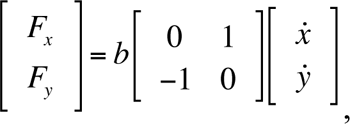

where *x*^ and *y*^ are hand velocity, and *F_x_* and *F_y_* are forces acting on the hand. Here, *b* was positive in the CW and negative in the CCW, and the magnitude of *b* was 20 N m^-1^s.

Each force field was associated with different audio-visual cues that were presented before reaching (Figure 1). Before reaching, in the CW, participants were presented with a red background, a red windmill-like diagram showing the direction and magnitude of rotational forces, and a high-frequency beep. Before reaching, in the CCW, they were presented with a blue background, a blue windmill-like diagram, and a low-frequency beep.

After 2-s cue presentation, one of the eight targets was randomly presented. After target presentation, participants were required to start within 1 s and reach the target within 225 ± 50 ms (time between exiting and entering start and target circles, respectively) using straight and uncorrected trajectories. The distance between the starting point and each target was 12.5 cm. The force field was off when participants returned to the start position and was on only during the recorded outward movements.

Visual feedback of hand position was suppressed during movements. The entire hand path was shown after movement termination. Participants were encouraged to learn a straight hand path with fixed movement duration by two types of rewards after each trial: one was given as a score of the current trial and the other as a score accumulated from the initial to current trial within that session. For each trial, participants gained 10 points for a successful start after target presentation (within 1 s), 20 points for a straight hand path when the hand was within 2 cm on the left or right of the straight line connecting the center of start and target circles, 30 points for stopping within the 0.15-cm radius target circle, and 40 points for movement duration within 225 ± 50 ms. A total of 100 points was awarded if a movement was fully successful.

### Procedure

The two force fields were presented either in blocks or randomly. In the block presentation, one cycle comprised eight consecutive trials in one of the two force fields, including randomly ordered movements to all eight targets. In the random presentation, two cycles consisted of 16 consecutive trials, including randomly ordered movements to all eight targets in the two force fields. In the random condition, a given force field was repeated no more than five times.

The BLOCK-BLOCK group (see Table 1 for Experimental protocol of each group) first learned two force fields in block presentation for four consecutive days as training and was then exposed to random presentation. On days 1 and 3, half of the eight participants performed one block comprising 112 movements (14 cycles) in CW. On day 2, they performed one block consisting of 14 cycles in CCW. On day 4, they performed one block consisting of 14 cycles in CCW and were then exposed to three blocks in the order of CCW, CW, and CCW, with each block comprising 64 movements (eight cycles). The other half was exposed to the force field in the reverse order. The BLOCK-RANDOM group first learned two force fields in block presentation for four consecutive days as training and was then exposed to random presentation. On days 1 and 3, 11 participants performed one block comprising 112 movements (14 cycles) in CW. On days 2 and 4, they performed one block consisting of 14 cycles in CCW. In the BLOCK-RANDOM group, the order was the same for all participants expecting that the similar results when the order was reversed since we find no significantly different behavior between the two orders in BLOCK-BLOCK group after subtracting the null field bias towards CCW. After block presentation on day 4, participants were exposed to 224 movements (28 cycles) in random order of CW and CCW as a test session.

RANDOM-RANDOM group first learned two force fields in random presentation for two consecutive days as a training session and were then exposed to random presentation as a test session. On days 1 and 2, 10 participants performed 224 movements (28 cycles) in random order of CW and CCW, and as a test session, they were exposed to 224 movements in random order of CW and CCW, interspersed with 32 catch trials without a force field. These catch trials were included to verify that participants were not employing a co-contraction strategy. The RANDOM-BLOCK group first learned two force fields in random presentation for two consecutive days as a training session and were then exposed to block presentation as a test session. On days 1 and 2, 12 participants performed 224 movements (28 cycles) in random order of CW and CCW. Then, half of the 12 participants were exposed to three blocks in the order of CCW, CW, and CCW, with each block comprising 64 movements (eight cycles). The other half was exposed to three blocks in the order of CW, CCW, and CW. We assigned 10 participants to a baseline control group (BLOCK-control group) who were exposed to three alternating blocks of 64 trials within one day as a baseline measure for reduction of interference by preceding training. Five of the participants in the BLOCK-control group were exposed to blocks in the order of CCW, CW, and CCW; the other five were exposed to blocks in the reverse order.

All groups except the BLOCK-control group experienced the same number of trials during the training session (224 movements for each force field). Prior to force field presentation, participants were familiarized with the apparatus and task during a block of 192 trials without any force fields (null force field: NF).

### Analysis

Adaptation to each force field was quantified by an error measure computed as the directional error (see above). To assess performance in a particular cycle, the median of the directional error of a set of eight movements in each force field within the cycle (or the two cycles for random presentation) was determined and averaged across participants for each cycle. Since null field performance was biased towards the CCW direction, we subtracted the median of the directional errors in the null field from that in the force fields for each participant. We compared the temporal evolution of these directional errors in the two force fields. For display reasons, the sign of the directional error was flipped for the participants who were exposed to blocks in the reverse order. To statistically confirm learning and retention, difference errors under CW and CCW were computed for each participant. The difference error was defined as the difference in directional errors between CW and CCW (CCW-CW), and indicated the average area enclosed by hand paths in CW and CCW.

### Effective learning rate

From equation 7, the effective learning rate for each module is given by *η_i_λ_i_* (*t*). The generalist responsibility *λ_generalist_* (*t*) increased as *σ* ^2^*specialist* increased when for a constant prediction error amplitude (see Supplementary Figure). Thus, the responsibility signal *λ_generalist_* (*t*) monotonously increased with the width of responsibility signal *σ* ^2^. A similar behavior can be shows for the specialist. Therefore, *σ* ^2^ can be used as an alternative to *λ* in estimating effective learning rate *η λ* (*t*). Then, (*η_generalist_ σ^2^_generalist_*)/(*η_specialist_ σ^2^_specialist_*) reflected the ratio of the effective learning rate between generalist and specialist modules.

### Statistical analyses

Statistical analysis was performed using Matlab statistics toolbox (The MathWorks, Inc.) and JMP software (SAS Institute Inc.). The p-value for significance was set 0.05.

## Acknowledgements

This work was supported by SCOPE, Ministry of Internal Affairs and Communications, the Strategic Research Program for Brain Sciences (SRPBS) JP17dm0107044, Ministry of Education, Culture, Sports, Science and Technology, JP18dm0307008, Japan Agency for Medical Research and Development (AMED), Funding Program for Next Generation World-Leading Researchers, and JSPS KAKENHI Grant Number 21H04425. NS was supported in part by grant NSF BCS 2216344.

## Notes

### Competing Interest Statement

The authors have declared no competing interest.

