## Supplementary Figure for "Inter-individual variability in motor learning due to differences in effective learning rates between generalist and specialist memory stores"

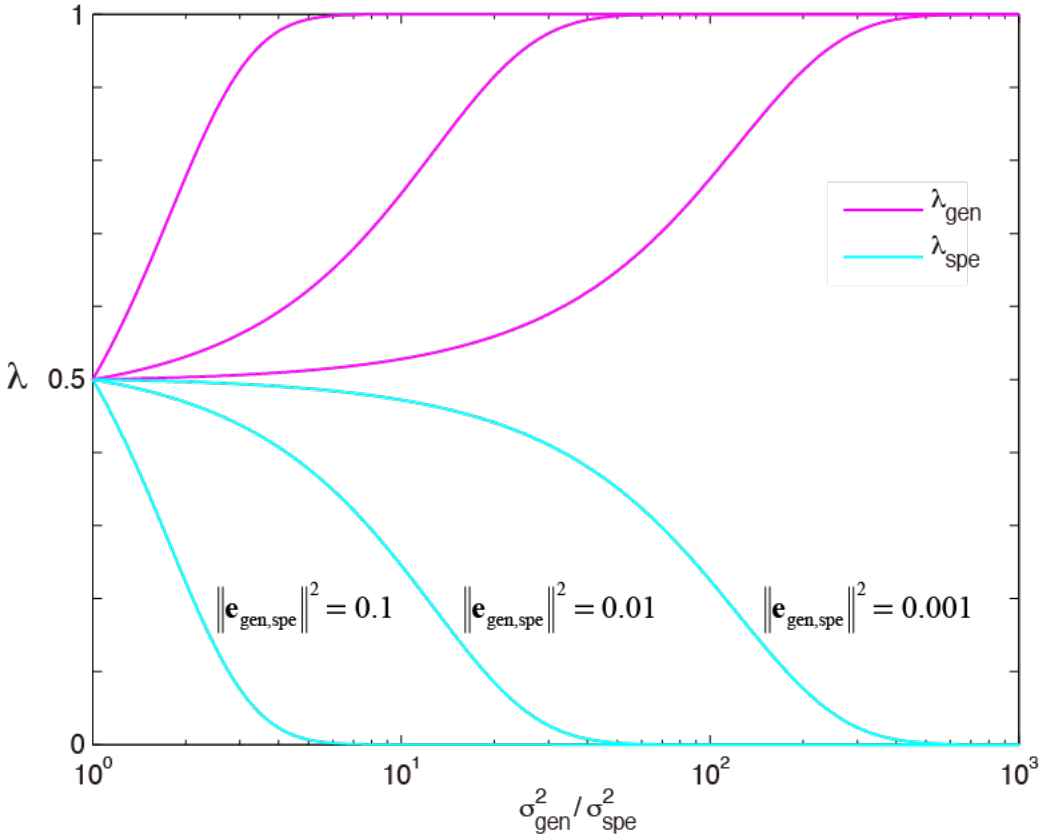


**Supplementary Figure:** The responsibility signal $\lambda_i(t)$ plotted against the ratio of the width of responsibility signal $\sigma^2_{generalist}/\sigma^2_{specialist}$. The magnitude of the prediction error $\|\mathbf{e}_i\|^2$ was fixed to 0.1, 0.01 or 0.001. $\lambda_{generalist}(t)$ monotonically increased as the $\sigma^2_{generalist}$ increased compared to $\sigma^2_{specialist}$ (magenta curves), while $\lambda_{specialist}(t)$ monotonically decreased (cyan curves).
